# A simple statistical framework for small sample studies

**DOI:** 10.1101/2023.09.19.558509

**Authors:** D. Samuel Schwarzkopf, Zien Huang

## Abstract

Most studies in psychology, neuroscience, and life science research make inferences about how strong an effect is on average in the population. Yet, many research questions could instead be answered by testing for the universality of the phenomenon under investigation. By using reliable experimental designs that maximise both sensitivity and specificity of individual experiments, each participant or subject can be treated as an independent replication. This approach is common in certain subfields. To date, there is however no formal approach for calculating the evidential value of such small sample studies and to define *a priori* evidence thresholds that must be met to draw meaningful conclusions. Here we present such a framework, based on the ratio of binomial probabilities between a model assuming the universality of the phenomenon versus the null hypothesis that any incidence of the effect is sporadic. We demonstrate the benefits of this approach, which permits strong conclusions from samples as small as 2-5 participants and the flexibility of sequential testing. This approach will enable researchers to preregister experimental designs based on small samples and thus enhance the utility and credibility of such studies.

## Introduction

Over the past decade, the fields of psychology and cognitive neuroscience have been embroiled in something of a methodological revolution, sparked by what is commonly referred to as the “replication crisis.” Large-scale efforts revealed that a considerable proportion of published research findings fail replication attempts, the corner stone of scientific inference (Open Science Collaboration, 2015). This has led to proposals for alternative statistical approaches, for instance based on the estimation of effect sizes (Cumming, 2014) or hypothesis tests using Bayesian statistics (Dienes, 2014; Rouder et al., 2009; Wagenmakers, 2007; Wagenmakers et al., 2018). It has also resulted in changes to the publication process, in particular the addition of peer review prior to data collection in the *Registered Report* format (Chambers, 2013; Chambers et al., 2015). Vetting and adjusting the experimental design and analytical approach prior to even collecting the first data point can control for biases and researcher degrees of freedom arising from methodological flexibility (Simmons et al., 2011). It is also simply good practice to set *a priori* thresholds for the evidence required to reach a conclusion.

These changes have thus far focussed primarily on the statistical framework of estimating population tendencies from limited samples. Such inferences are necessary for answering many research questions, such as quantifying the effect of a medical treatment or a psychological manipulation. However, this often goes together with requiring ever larger sample sizes to test increasingly more subtle effects. When used unwisely, this approach is wasteful and could prove unsustainable. There are only limited resources for scientific research; expecting vast samples (possibly collected by multi-site consortia) necessitates diverting funds that could otherwise have been shared by several teams to address a range of questions. The inevitable consequence is that scientific progress will slow down. However, contrary to the ingrained belief of many journal editors and reviewers, group studies based on large sample sizes are not necessary for all kinds of research (Smith & Little, 2018). Indeed, for some research the mindless use of group statistics (Gigerenzer, 2004) can even hinder the interpretation of results (Ince et al., 2022).

The purpose of psychology and neuroscience research is to discover the mechanisms of the mind and brain. To answer many research questions, the average population effect is at best uninformative, or even irrelevant. Sensory psychophysics studies regularly employ a small sample (small-N) approach in which hypotheses are tested within each individual observer. This considers each observer as an independent replication. Some studies can demonstrate convincing results with a sample as small as two subjects^1^. Rather than inferring the average effect in the population, these studies ask whether the phenomenon in question is *universal* to human perception. Provided the individual tests are reliable, demonstrating their existence in a small number of people is ample evidence. Importantly, while this approach is common in psychophysical studies and single-cell recordings in primates, it should also be suitable in many other scenarios, such as neuroimaging, human electrophysiology, and psychophysiological experiments. Critically, it is also readily applicable to many classical experimental psychology studies that typically use the classical group statistics framework.

This requires making a crucial consideration: is our aim to discover knowledge about fundamental mechanisms or processes, or is it instead necessary to estimate the magnitude of a phenomenon? The latter aim tacitly assumes that individual differences across the sample reflect genuine deviations from the average. Body height can be measured with great precision; armed with a standard tape measure, even a single measurement should have minimal error. The variability in height across the sample therefore reflects how much true variation there is between people. This scenario calls for classical group statistics. In contrast, the same cannot be said for most studies in experimental psychology. A task measuring variables like reaction time, the proportion of correct trials, or a match of perceptual brightness will depend crucially on the number of replicate trials conducted in each individual subject. Too few trials result in differences between individuals that do not reflect the true population variance, but rather the variability contaminating the individual estimates. It makes far more sense to collect as many trials per subject as feasible, and thus establish separately if the effect manifests in each person. There may still be genuine individual differences, but any *fundamental* feature of the human mind should be detectable in everyone – except for rare instances when the test fails.

But how many subjects are necessary to confirm the existence of a phenomenon? And how does this depend on the reliability of individual tests? Without concrete answers to these questions, small sample studies will continue to run afoul of ill-advised editorial decisions and reviewer recommendations. What is worse, the current climate fosters widespread misconceptions about what makes good science.

### Binomial model comparison

Here we present a basic framework for estimating the statistical evidence about results from small sample designs. We call this the ‘small-N test’ or ‘universality test’. The underlying logic is that if an effect either holds for virtually all cases (except for one or two exceptional ones) or for practically none, then even a very small sample can reveal whether it is a universal feature. Consider the classical Stroop effect (Stroop, 1935). People typically suffer a reaction time cost for reading the word “red” written in another font colour. Provided each subject completes a sufficient large number of trials, we would expect every healthy, neurotypical person to show this effect, if it is indeed a universal feature of human cognition. The few who do not are probably statistical exceptions – this can happen simply by coincidence but also due to nuisance factors, such as whether the subject’s attention was diverted (likely in an online experiment) or because, unbeknownst to the experimenter, the person participated in the experiment while under the influence. In contrast, we would not expect an individual subject to predict the outcome of 100 coinflips significantly more than 50 times. There may be a small number of subjects who score higher, but these are probably rare flukes. This result would certainly not support the notion that precognition is a universal human ability.

In statistical terms, our approach compares the likelihood of two opposing models: the alternative (experimental) hypothesis, H_1_, assumes that the phenomenon of interest is a universal feature across all subjects. Any failure to detect it in a subset of individual tests should merely be due to lacking sensitivity of these tests, perhaps because of lapses in performance or other individual differences skewing the measurement. In contrast, the null hypothesis, H_0_, assumes that the phenomenon is not universal, and that any individuals showing a significant effect are either rare outliers or random flukes.

The model comparison is then conducted via a likelihood ratio, similar to what is commonly done for statistical inference via Bayes Factors or likelihood tests (Jeffreys, 1961; Rouder et al., 2009; Wagenmakers, 2007). To determine the likelihood of H_1_ we calculate *P(D|H_1_)*, the binomial probability density of obtaining the observed number of significant subjects, *k*, out of *n* subjects in total, given the sensitivity of the individual test. The sensitivity is 1-β, where β is the false negative rate. The likelihood of H_0_ is *P(D|H_0_)*, the binomial probability of observing the same result (*k* successful out of *n* tests in total) under the assumption that the effect occurred by chance. The chance probability of the individual tests is their significance threshold, or the false positive rate α. To illustrate, Figure 1 plots two example binomial probability density functions for a set of 7 subjects out of whom either only 1 or 5 yielded significant findings. The evidence ratio, which we labelled ε to indicate it is in the context of the small-N framework, is then given by,

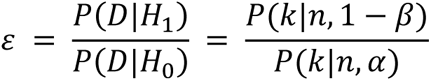

**Figure 1.**
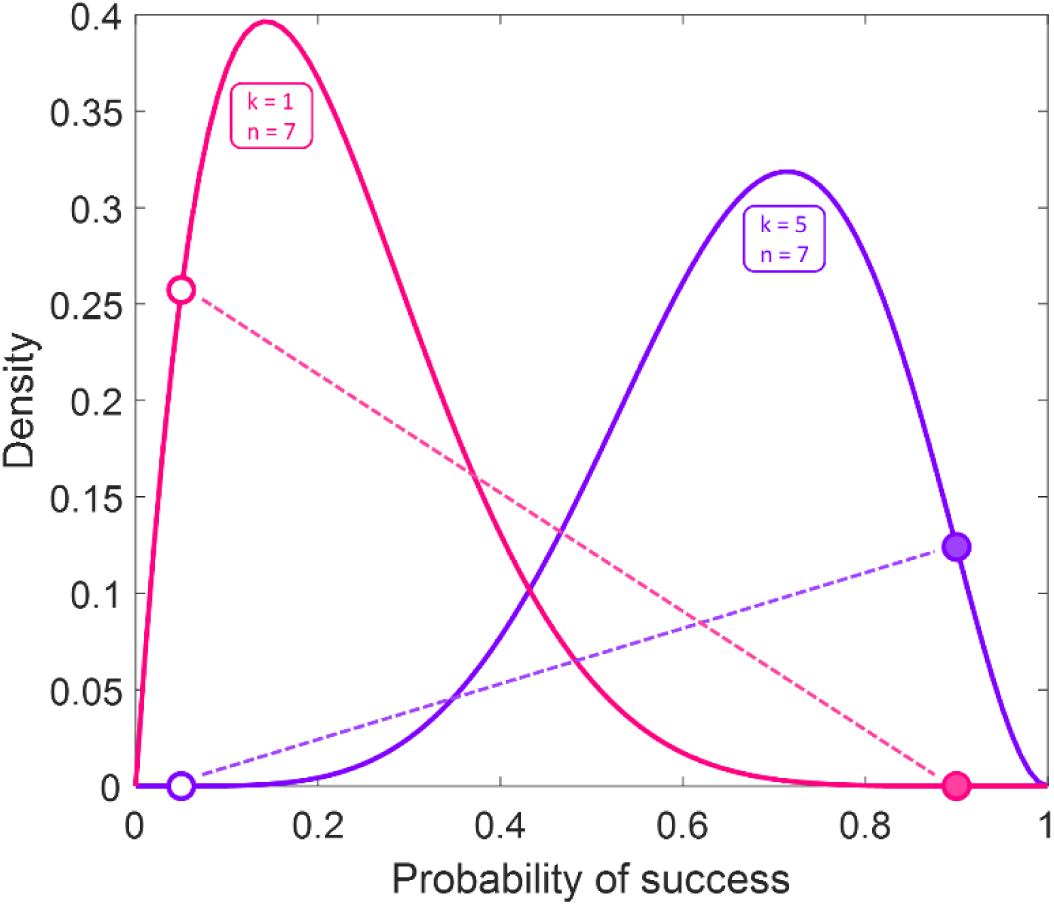
Binomial probability density functions for achieving k=1 success (pink) or k=5 successes (purple) out of n=7 total tests (subjects) plotted against the success probability. Filled discs denote the binomial probability when success probability is 0.9, a common assumption for sensitivity. This is the likelihood of the alternative hypothesis, *P(D|H_1_)*. Open discs denote the binomial probability when assuming successful tests occur by chance, the false positive rate 0.05 of the individual tests. This is the likelihood of the null hypothesis, *P(D|H_0_)*. The evidence ratio, ε, is the ratio of these likelihoods. The ratio is large/small when the alternative hypothesis is more/less likely than the null hypothesis (visualised by dashed lines).

This quantifies how much more likely the observed result D is under H_1_ than under H_0_. In pragmatic terms, if ε is high we would expect the result to replicate when running another subject. Since the calculation of the evidence relies on the error rates, α and β, this necessitates an experimental design that can minimize these errors. Determining the false positive rate is relatively straightforward, especially in designs that use standard significance tests at the level of individual subjects, such as through a p-value or a confidence interval. Deriving the false negative rate is more complicated, especially as sensitivity can vary between subjects.

To illustrate, let us assume perfect sensitivity. If the phenomenon is universal, the test will always detect it. If the test also had perfect specificity, α=0, we would only need to test a single subject to draw a conclusion about whether the phenomenon exists. Such a perfect test is rare, if it exists at all; even highly reliable procedures will carry at least a minor risk of false positives. Thus, assuming a conventional significance threshold of α=0.05 for the individual tests, after running a single subject we can obtain the following evidence ratios: if this single test fails to reach significance, this rules out the universality of the effect because with perfect sensitivity we should have confirmed H_1_. However, if the individual test is significant, this indicates that H_1_ is 20 times more likely than H_0_. Such an evidence ratio is relatively strong, although maybe not sufficiently compelling to stop the experiment after only a single subject.

What about running two subjects? If both test significant, the evidence is ε=400 (=0.05^-2^), which seems like a convincing result. But because the sensitivity of the individual test is perfect, any other result would support H_0_, however, and completely rule out that the phenomenon is universal. Naturally, perfect sensitivity is just as improbable as perfect specificity. Even the most reliable test sometimes fails to detect a phenomenon. Statistically, this means we might assume a sensitivity somewhat less than perfect, perhaps 0.95. Now, if we test only two subjects and find a significant effect in both, this produces an evidence ratio of ε=361, the ratio of *P(2|2, p=0.95)* and *P(2|2, p=0.05)*, or 0.95^2^/0.05^2^. Conversely, finding no significant effects, does not rule out its existence but gives the inverse ε=1/361, approximately 0.0028. These are both still very conclusive results for H_1_ or H_0_, respectively. Crucially, finding only one of the two subjects was significant means that the result is completely inconclusive (ε=1). In this case, more subjects are required to make an inference. If we tested a third subject, and their result was also significant, this would again produce a moderate ε=19. Three significant out of four subjects would bring back compelling evidence at ε=361.

What this toy example demonstrates is that the small-N approach naturally lends itself to sequential testing. This is unsurprising, given that the logic is essentially the same as for inference using Bayes Factors, which is commonly used for sequential sampling in group studies (Rouder, 2014). Crucially, this means that researchers can (and should) define a threshold for the evidence ratio needed to make an inference. The choice of threshold is up to researchers and depends on the nature of the study. Confirming a well-known sensory aftereffect is probably compelling with ε≥16 (i.e., corresponding to a conventional significance threshold of 0.05 and sensitivity of 0.8). In contrast, testing extraordinary claims of precognitive abilities, as in a study that was partly responsible for triggering the replication crisis in psychology (Bem, 2011), should probably demand more extraordinary evidence. As a rule of thumb, we chose a relatively conservative threshold of 50 (or 1/50=0.02, respectively).

### Determining the sensitivity

One problem common to experimental design, not only for this small-N approach but also classical group studies, is how to determine the sensitivity, or the statistical power, of the experiment. Calculating power requires a predefined effect size of interest. This is often based on the effect sizes reported in previous research, although these may not always seem appropriate because previous studies used different experimental paradigms or different dependent variables. Most previous research is also affected by publication bias, irrespective of the methodological rigour or controls put in place (Kühberger et al., 2014; Masicampo & Lalande, 2012; Newcombe, 1987; Rosenthal, 1979). There may be an unknown number of attempts at similar experiments whose results were not strong enough to make it into the literature. This means that predicted effect sizes based on previous research are likely inflated. A more principled approach could be to determine an effect size of interest based on the practical significance of an effect.

The latter type of consideration is particularly crucial for the small-N approach. In most contexts, it should be possible to design the tests in individual subjects such that they are highly sensitive to reveal practically meaningful effects. For instance, when estimating an illusion magnitude (subjective perceptual bias) by fitting a psychometric curve, a practical consideration could be that the detectable bias should exceed the just noticeable difference (objective discrimination ability). The experiment should therefore collect enough trials to ensure that such a bias can be detected reliably. Similar considerations could determine the sensitivity of neuroimaging studies, e.g., the correlation between voxel response patterns from a brain region within a single subject. Power analysis can determine the minimum correlation detectable with the number of voxels in a brain region. Critically, because sensitivity varies between subjects, this estimate should aim to maximize sensitivity for the weakest link, that is the subject thought to have the lowest sensitivity, due to the smallest number of voxels, trials, cells, etc.

However, even in the absence of a clear estimate of individual sensitivity, it is possible to estimate the evidence ratio by plotting the full range of possible sensitivities, i.e., between the false positive rate α and 1 (we naturally assume that in well-designed studies, sensitivity should always exceed α). Evidence curves for a range of typical sample sizes (Figure 2) show that when the evidence favours the null hypothesis, that is, when only a few subjects test significant, the support for H_0_ increases steadily with sensitivity. In contrast, when most subjects in a sample test significant, the evidence peaks at an intermediate sensitivity and then declines. As already discussed, unless all subjects test significant, the evidence ratio with a perfect sensitivity of 1 must be 0. Therefore, highly sensitive tests also imply a high likelihood that all subjects test significant and any divergence from this result provides evidence *against* H_1_. The peak of the curve in fact corresponds to the observed proportion of significant findings: e.g., with 2 significant out of 3 subjects the evidence curve peaks at ∼0.667 sensitivity.

**Figure 2.**
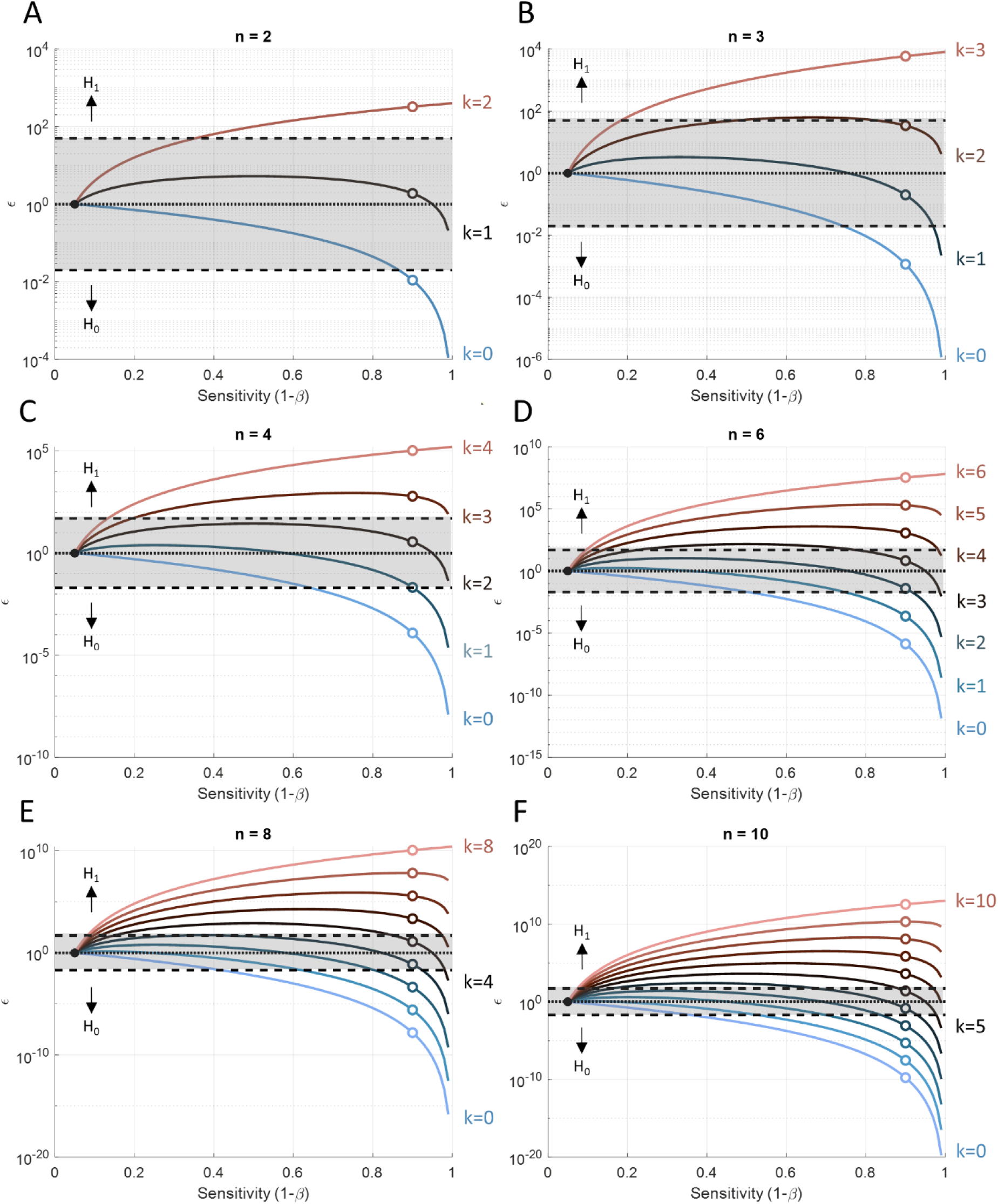
Statistical evidence plotted as a function of the sensitivity of individual tests. Each curve denotes the evidence ratio ε obtained for a given number of significant tests in the sample, k, ranging from no significant tests (k=0, blue) to all tests significant (pink). Curves are shown for sample size n=2 (A), n=3 (B), n=4 (C), n=6 (D), n=8 (E), and n=10 (F). The dashed black lines and grey shaded areas indicate the region of inconclusive evidence where 0.02<ε<50. Evidence ratios above or below this region indicate compelling support for H_1_ or H_0_, respectively. For reference, open circle symbols indicate sensitivity of 0.9.

Inspection of these evidence curves allows an insight into the overall evidence for H_1_, even when the sensitivity is uncertain. By determining the range of likely sensitivities of the individual tests one can identify the strength of the evidence. If a researcher is confident that their individual tests have near-perfect sensitivity, then any result but an overwhelming majority of subjects testing significant argues against the universality of the phenomenon.

For most experiments, such extreme sensitivity is however unachievable; therefore, a fraction of non-significant subjects can still provide strong evidence for H_1_. With a sample size of n=2 (Figure 2A), if both subjects test significant this would pass the ε>50 criterion at a relatively low sensitivity of only ∼0.354. However, if no subject tests significant, this would not reach the evidence threshold for H_0_ of ε<0.02, unless sensitivity was at least ∼0.866. Clearly, if the researcher has high confidence in the individual tests, such a result convincingly argues against the universality of the phenomenon. Finally, if only one of the two subjects tests significant, the evidence curve always stays between 0.02 and 50. More data must therefore be collected regardless of the sensitivity of individual tests.

The relationship is similar for larger sample sizes (Figure 2B-F). Larger samples simply have the consequence of shifting down the required sensitivity to reach conclusive evidence for either hypothesis. Just as is the case for classical group statistics, it is easier to obtain evidence supporting the existence of an effect than it is to support its absence. At n=10, the necessary sensitivity for obtaining ε>50 if all subjects test significant is ∼0.074. This would constitute an overwhelming false negative error rate and suggests the need for a better experimental design. Conversely, if none of the ten subjects test significant, the sensitivity needed to obtain support for H_0_ at ε<0.02 is ∼0.357, still a relatively insensitive test. This example underlines the fact that larger samples are not superior in this approach; rather they only benefit poorly sensitive studies.

We summarise this relationship in Figure 3. This shows across a range of sample sizes the sensitivity required for passing the evidence thresholds for H_1_ and H_0_, respectively, when either all or none of the subjects are significant. As the name implies, the small-N approach is designed to provide conclusive evidence for small samples. It demands maximising the sensitivity of individual tests. If high sensitivity can be achieved, a conclusive study can be conducted with substantially fewer than ten subjects. However, if sensitivity cannot be optimised, the approach is ill-advised, and a conventional group average design is more suitable.

**Figure 3.**
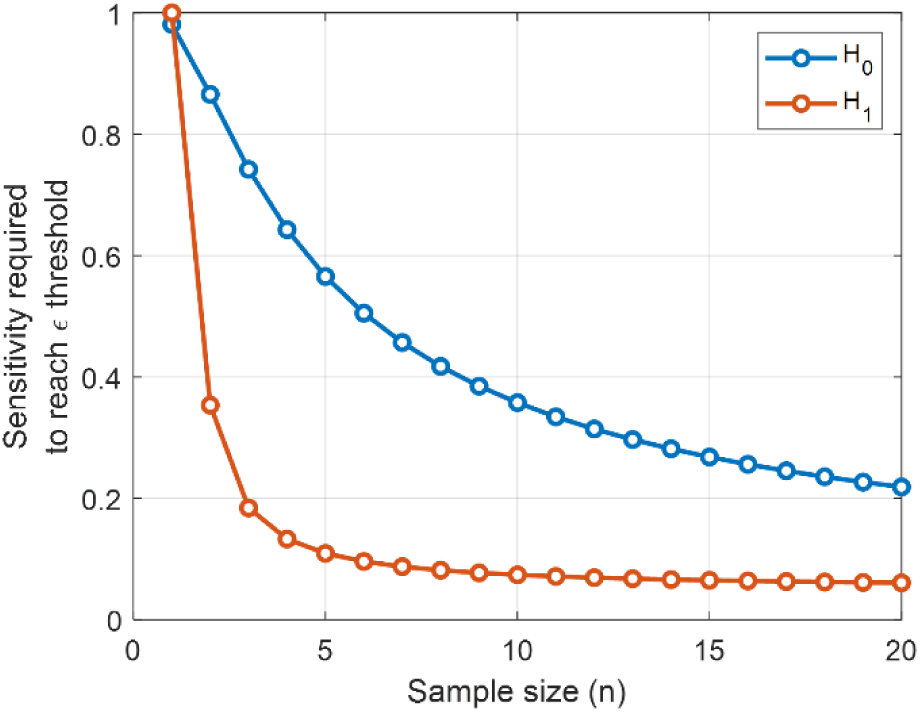
Sensitivity necessary for obtaining conclusive evidence with a given sample size. Curves plot the required sensitivity needed for ε>50 for supporting H_1_ when all subjects test significant (red) or ε<0.02 for supporting H_0_ when no subject tests significant (blue).

### Comparison with group statistics

By now, it has probably become apparent that the small-N approach is particularly suitable for robust effects. If the phenomenon manifests in most of the sample, this strongly supports its existence. Naturally, a highly consistent effect across the whole sample is also likely to be significant in a classical group test. Why then should we use the small-N approach instead of simply using classical group statistics on the small sample? One reason is that many investigators regard significant findings from small samples with suspicion, even if the results are compelling. More importantly, as we discussed, the small-N approach asks a different question than group statistics. The small-N approach is indicated whenever we make inferences about universal features common to all subjects rather than quantifying the magnitude of this feature. To conclude that typically-developed humans are born with an appendix, we need only confirm they have this vestigial organ; we would not test that the average human appendix is longer than 0 cm.

To illustrate the superiority of the small-N approach for such research questions, we sought to replicate a well-established effect from perceptual psychophysics, known as the oblique effect (Appelle, 1972; Furmanski & Engel, 2000; Vogels & Orban, 1985): observers’ ability to discriminate the orientation of oblique orientations (that is, near 45° or 135°) is considerably worse than performance near cardinal orientations (0° and 90°, corresponding to horizontal and vertical). This difference in performance is thought to reflect differences in the neural populations in visual cortex coding for the different orientations (Coppola, White, et al., 1998; Furmanski & Engel, 2000), possibly related to the statistical distribution of oriented edges in the natural environment (Coppola, Purves, et al., 1998). We are inclined to treat this is a fact about human visual performance.

We collected orientation discrimination data for the oblique and vertical orientations in a sample of n=17 observers (see Supplementary Information). We quantified this using a psychometric curve fit where the bandwidth is an indication of discrimination ability: narrower bandwidths correspond to better performance. This confirmed our prediction that average bandwidth for judging orientations near 90° was smaller than near 45° obliques (one-tailed t-test for μ>0, t_16_=13.88, p<0.0001; Bayes factor with a default Cauchy prior scaled with r=√2/2, BF_10_= 74,700,000), an example of a robust effect. Conversely, comparison of the two oblique orientations revealed no significant difference in performance (t_16_=-1.14, p=0.8644) and Bayesian inference favours the null hypothesis that bandwidths near 45° were not greater than near 135° (BF_10_=0.13). We use this dataset as an example of a compelling null result.

What would have happened had we analysed this experiment using the small-N approach? Our psychometric curve fit also allowed us to make statistical inferences for each individual observer. Doing so revealed that the oblique effect (bandwidths near 45°>90°) was significant at p<0.05 in all but one observer. Even this non-significant observer replicated the effect numerically. Under the assumption that this phenomenon is a replicable effect that should manifest in all observers, we take this as an estimate of the sensitivity of our test, 0.941. In contrast, the null effect (bandwidths near 45°>135°) was only significant in one observer. This corresponds to ∼0.059, close to the nominal false positive rate expected of the test.

Thus, using α=0.05 and a sensitivity of 1-β=0.941, we calculated the evidence ratio when incrementing the sample size sequentially in the order individuals were tested (Figure 4A, purple curve). This showed that even with two observers, the evidence for the oblique effect already passed the threshold of ε>50. Testing more observers only further increased the evidence. By the time we reached observer 12, whose results failed to reach significance, this could hardly dent the evidence. Importantly, for the null effect we also reached conclusive evidence in favour of H_0_ with two subjects already (Figure 4B, purple curve). By the time, we ran observer 15, who tested significant despite expecting no difference, the evidence for the null hypothesis was already overwhelming. Essentially, using the small-N approach we could have ended data collection after two observers already – unheard of in conventional group studies.

**Figure 4.**
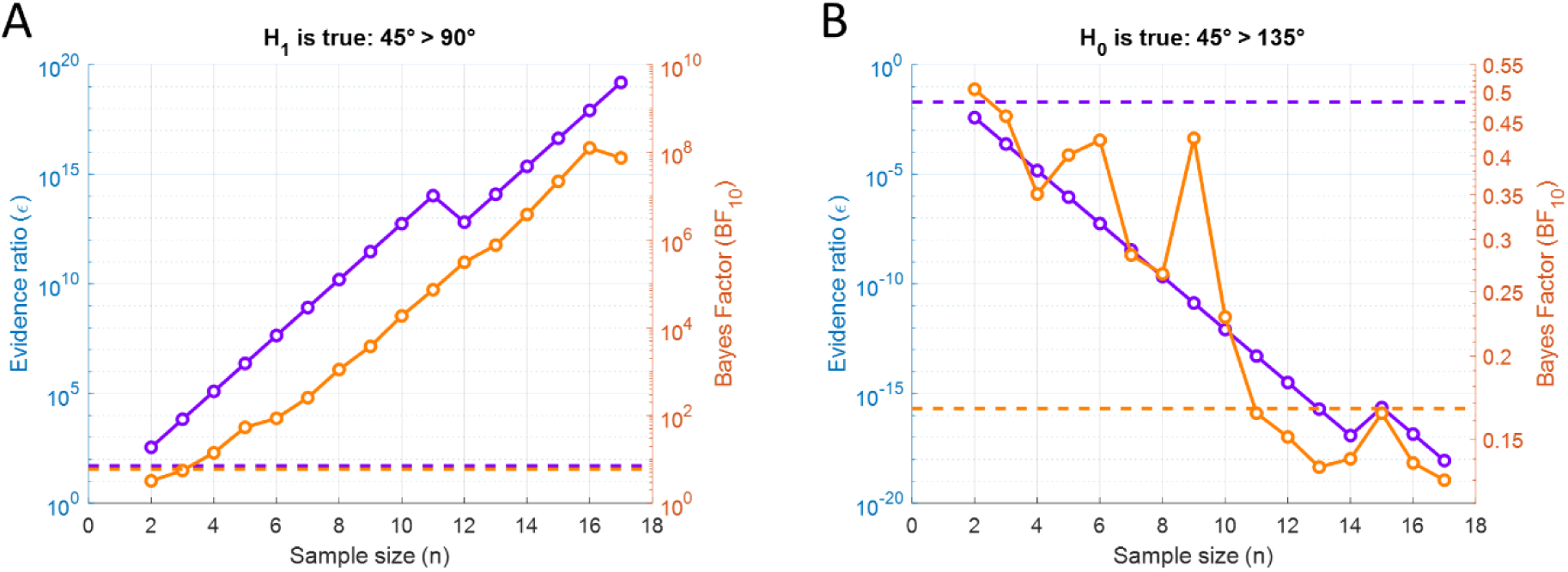
Statistical evidence in oblique effect experiment as a function of sample size. Evidence ratio ε (purple) of the small-N test, and Bayes Factor BF_10_ of a one-sample t-test for μ>0 (orange), were plotted against the sequentially increasing sample size, shown separately for the assumed true effect (A) and the assumed null effect (B). Note separate y-axes for the two measures. Dashed horizontal lines denote the thresholds for conclusive evidence supporting H_1_ (ε>50, BF_10_>6) or H_0_ (ε<1/50, BF_10_<1/6), respectively.

In comparison, incrementing sample sizes using classical group statistics would have forced us to continue the experiment (Figure 4, orange curves). We only report Bayes Factors for this analysis because sequential testing skews frequentist statistics (Rouder, 2014; Simmons et al., 2011), although adjustment procedures exist for this situation (Lakens, 2014). For the oblique effect (Figure 4A), after 3 observers we would have obtained a Bayes Factor above the criterion of BF_10_>6 used as evidence threshold in the *Registered Reports* format by many journals (e.g., *Cortex:* https://rr.peercommunityin.org/about/pci_rr_friendly_journals#h_4920688494031618419330727). This would therefore have entailed testing one more observer than the small-N approach. This may not seem like a big difference; however, even with a strong Bayes Factor like this (BF_10_=14.1), experience tells us that many people would find this result unconvincing. In fact, we wager that this would lead to immediate desk rejection by the handling editor at some journals. On the other hand, the small-N approach demonstrates that even two observers testing significant is already extremely compelling evidence.

However, the true benefit of the small-N approach becomes clear for the null effect. The group analysis only produced any evidence in favour of the null hypothesis (BF_10_<1/6) by the time we had tested observer 11. We would have needed to collect more than five times the sample size required for reaching the same conclusion using the small-N approach. The Bayes factors for the null effect were also highly unstable, jumping up and down as the sample size increased. After testing observer 15, whose results showed an unexpected significant difference at the individual level, the Bayes factor came dangerously close to falling above the criterion for H_0_ (BF_10_=0.164). With small samples, individual effect sizes can have a profound effect on the sample mean, resulting in considerable fluctuations in the evidence of group tests. In the small-N approach, however, the evidence for H_0_ steadily increased, with observer 15 causing only a tiny blip. Note also that these conclusions do not strongly depend on the chosen evidence threshold. The situation would be similar for a criterion of BF_10_>3 or BF_10_<1/3, respectively, especially considering that our criterion for the small-N approach of ε>50 or ε<0.02 could be considered highly conservative.

One caveat is that the small-N approach depends on the sensitivity of individual tests. In our example, we estimated sensitivity from the assumed false negative rate. We could only do so because he had collected a larger sample, and because we were already convinced that the oblique effect is universal. In typical experiments, however, one would need to rely on other estimates. For a psychophysical study such as this, sensitivity is usually high, provided enough trials are collected per observer. Through simulations one could determine the required number of trials for estimating the perceptual effect, although this can be a relatively complex because psychophysical performance depends on various factors. However, as already explained, even without precise knowledge of the achieved sensitivity, inspection of the evidence curves (Figure 2A) permits the conclusion that two out of two observers showing a significant oblique effect strongly support its existence; the evidence is already conclusive even with comparably low sensitivity (Figure 3, red curve). For supporting the null effect, much higher sensitivity would be required (Figure 3, blue curve), so we can only draw this conclusion with highly sensitive tests. To support a null result, one would likely need to collect a few more observers, around 4 or 5 in total. However, the required number would still be considerably smaller than the 11 we needed using classical group statistics.

### Examples from the literature

We suspect most researchers will consider samples of only two subjects to be very small. However, there are examples of this in the literature. A seminal psychophysical study (Field et al., 1993) used only two observers – who also happened to be authors of the article – to characterise the parameters that govern performance in the contour integration (‘pathfinder’) task where subjects must detect the presence of a group of aligned Gabor patches in a cluttered field of randomly oriented distractors. Despite the small sample of non-naïve observers, the results are clear and compelling. It would be difficult to estimate the sensitivity of these tests, but we can be confident that it far exceeds what is necessary to detect the reported effects. The study reported five experiments, four of which show the same significant effect in both observers and the fifth shows a convincing null result. While not commonly used in other subfields of experimental psychology beyond perceptual psychophysics, there is no reason why this approach cannot also be used for other phenomena, such as distractor interference, executive control, memory tests, or stimulus-response priming. The key is simply to collect sufficient data in each subject to detect the effect at the individual level.

There are also good examples from the neuroimaging literature. Another famous study used an encoding model to demonstrate that the identity of natural images can be accurately predicted based on response patterns from early visual cortex (Kay et al., 2008). They demonstrated this result in two subjects. Nobody would doubt this result, given the convincing reliability of these effects within each subject. While this may be an extreme example, the same concept applies to other neuroscience findings whenever many data points are used to make an inference at the level of individual subjects, for example when comparing the response of a set of independently localised voxels, a population of neurons in single-cell recordings, or a representational similarity analysis using patterns of responses. We also recently published neuroimaging studies with small sample sizes, but where the result for each subject is an activation map comprising hundreds to thousands of data points (Huang et al., 2023; Urale et al., 2023). This research was completed prior to the development of our present framework. Instead, we set a “consistency criterion” specifying that most individual subjects (5 of 7 and 6 of 8, respectively) must show a significant effect to consider the overall result meaningful. The results are compelling but clearly the criterion is arbitrary and its interpretation not intuitive. Using our small-N approach, we can now easily determine the evidence ratio for these studies. The evidence strongly favours the hypothesis that the effects are universal.

A corollary is that methods like representational similarity analysis (Kriegeskorte et al., 2008) are more powerful than frequently used multivariate classification decoders (Cox & Savoy, 2003; Haynes & Rees, 2005; Kamitani & Tong, 2005). The latter typically predict condition labels (stimulus categories, mental states) based on patterns of brain activity on a trial-by-trial basis. The sensitivity of this analysis therefore depends solely on the number of trials producing the classification accuracy for each subject. This number is typically small, and so the sensitivity is also low. In contrast, a representational similarity analysis correlates the voxel response patterns of independent stimulus conditions. The sensitivity is given by power analysis for correlations with the number of voxels in the pattern, based on classical power calculations. Using the small-N approach, a significant correlation in 2-3 subjects could already be compelling evidence, whereas classification accuracy in a typical design is unlikely to be significantly greater than chance in all three subjects.

Finally, the small-N approach can also provide meaningful answers where the outcome of conventional group statistics can be misleading. Claims of precognitive abilities in classical psychology experiments were supported by significant group statistics (Bem, 2011). Specifically, experiment 1 of this study reported performance for guessing the outcome of future trials with 53.1%, significantly above the chance level of 50% (t_99_=2.51, p=0.01). The experiment used 100 subjects, each of whom performed 18 or 12 trials, respectively, in the critical condition. The average precognitive ability therefore constitutes less than a single trial above chance. The author’s argument was that to detect a phenomenon as subtle as this requires averaging performance over many subjects.

However, by instead increasing the number of trials performed by each subject, the individual sensitivity should be sufficient to detect extremely subtle effects, even smaller than the reported finding. Thus, the same experiment could have been conducted using the small-N approach. Admittedly, this would entail running thousands of trials in each person. But the evidence of finding significant above-chance performance in only 3-4 subjects with a high-powered procedure would be far harder for sceptics to ignore than a significant group effect in a sample of 100 subjects. On this note, a recent replication attempt of this experiment used a methodologically highly rigorous procedure and found no evidence for precognition (Kekecs et al., 2023). This was based on a sample of n=2220 subjects. It could be more efficient to test four people 200 times, than running an experiment on such a huge sample; but of course, there could be reasons why only a small number of trials can be collected per subject; for instance, some experiments are specifically designed with one critical trial in mind (Dijkstra & Fleming, 2023).

## Discussion

We introduced a simple framework for estimating the statistical evidence of studies using small samples, typically comprising fewer than ten subjects. Despite justified concerns about the statistical power and replicability of research in psychology, neuroscience, and other areas of life sciences, obviously not all studies in those fields require large sample sizes. Any experiment that can provide reliable data from individual subjects, based on large numbers of trials or observations, is suitable for small-N analysis. Rather than estimating the average population effect, the small-N approach treats each subject as an independent replication. Most importantly, it does not require extensive knowledge of complex statistical methods because it is simply based on the binomial probability function. This can be calculated using most standard statistical analysis packages or even online calculators. We also provide MATLAB and R code and an easy-to-use Excel spreadsheet with this article for the readers’ convenience (Schwarzkopf & Huang, 2023a).

Small sample studies are commonplace in several subfields of neuroscience research, ranging from psychophysical studies to neurophysiological recordings in animal models. While this has been pointed out previously (Smith & Little, 2018), to date no straightforward frameworks for interpreting the evidential value of such studies have been described. By formalising this approach, we hope to expand the use of small sample approaches to other research contexts where the benefits of this approach have thus far not been realised. For instance, it should be readily applicable to many experimental psychology paradigms, such as visual search tasks, memory tests, distractor-interference, to name but a few. It can be applied in human electrophysiological, neuroimaging, and brain stimulation experiments, and psychophysiological measurements like pupillometry, eye tracking, or galvanic skin responses. The key is that each subject must be tested with enough replicate trials/data points to achieve high sensitivity at the individual level. In turn, this prescribes a within-subject design; all experimental conditions should be tested in each subject. We hope to raise awareness among researchers (including journal editors and reviewers) that such an approach can yield robust and meaningful results.

Our empirical example, using the well-established oblique effect (Appelle, 1972; Furmanski & Engel, 2000), shows that the small-N approach produced conclusive results with only two subjects, while conventional group analysis would have required more than five times this number to support a null result. Consider the waste of resources, work hours, and funds that this might entail in a real experiment. Human volunteers are precious. Running unnecessary experiments is counterproductive: many subjects who might be disinclined to return for more experiments could have participated in other studies. It is also unethical, and this issue becomes even greater in animal studies, where researchers have an obligation to reduce the number of subjects and minimise their suffering.

Depending on the situation, the small-N approach also allows us to gain insights into scenarios with individual differences. For example, our empirical demonstration was based on the rationale that the oblique effect manifests in all subjects. We found a single observer without a significant result. This is most likely a false negative; while sensitivity was presumably high, there is a small chance that one test will fail to reach significance. The evidence ratio shows this result is far more likely assuming the effect’s universality than under the null hypothesis of random false positives. However, it is also possible that the effect is common but *not* universal. Repeating the experiment on this observer could put this hypothesis to the test: if the effect is universal, it is highly likely significant on the second test. However, if the effect were due to true individual differences, it would be more likely that it again fails to reach significance.

### Relationship to prevalence

In this context, we note that a similar approach to ours has recently been described which uses a Bayesian framework to estimate the prevalence of a phenomenon in the population (Ince et al., 2021; Ince et al., 2022). This approach complements our small-N approach but there are a few points of divergence. First, as the name implies the prevalence approach quantifies the frequency of the phenomenon, and this could be relatively low. The prevalence approach could therefore be useful for revealing individual differences or significant outliers (Ince et al., 2022). In contrast, our small-N approach assumes that an effect is universal (under the alternative hypothesis), or that any individual incidence of it either arose by chance, due to nuisance factors, or because it is rare (null hypothesis). A corollary is that the prevalence approach may still require relatively larger samples to draw confident conclusions about prevalence. Conversely, the small-N framework is literally designed to minimise the number of required subjects.

Caution is advised when interpreting evidence for a universal effect. Imagine an alien visitor to our planet who sets out to test the hypothesis that all human beings have red hair. Given the low prevalence of red hair, it would be extremely unlikely to find that 4 out of 5 randomly selected individuals are redheads. Thus, the evidence ratio will almost certainly reject the hypothesis that red hair is universal. But now consider the opposite scenario. Globally, dark hair is highly prevalent. Thus, it is likely that a majority of five people the alien randomly selected will have dark hair. Taken at face value, the small-N test would incorrectly suggest that all people have dark hair (i.e., prevalence of 100%).

This example highlights the importance of considering the plausibility of the hypothesis. How realistic is it that all people are dark-haired? Prior knowledge can be part of the equation here. While we only use five random individuals to test the hypothesis formally, unlike the alien visitor, most human researchers have prior experience of meeting people with other hair colours. More importantly, the hypothesis can also be informed by biological evidence. Certain genes can determine hair colour and some of these are recessive. This enables an informed guess as to the distribution of hair colours. The alien may not have access to this information, and it would therefore behove them to reserve judgement. They *could* however already conclude that dark hair is widespread.

The same caveat applies to our empirical example of the oblique effect. Based on a small sample of 2-3 observers, we cannot rule out the possibility that a small proportion of people show no oblique effect. We however hypothesise that it is universal because this is more biologically plausible than people varying considerably in the functional architecture of their visual system. Nevertheless, this is a possibility that would require further study, assuming there is any reason to suspect it. The prevalence statistics mentioned above can also help adjudicate in this situation. Note, however, that this is not unique to the small-N approach, but similar caveats apply to *all* statistical models: just because we find a significant linear regression between two variables does not necessarily mean that a non-linear relationship is not an even better model.

It should be noted that when an effect is relatively common but not all subjects in the sample test significant (e.g., 2 out of 4), this will often result in an inconclusive evidence ratio (Figure 2C). Ideally, this should be addressed by collecting more data. Assuming that sensitivity has been maximised, if the evidence does not become conclusive (supporting either H_1_ or H_0_), this indicates that the effect is not universal, but that it did not arise by chance either – therefore, it is a relatively prevalent effect.

### When small samples are appropriate

Importantly, the small-N approach is not a replacement for conventional group statistics. Rather it adds to the toolbox of scientific research when the research question is suitable. Many research questions could have been better answered using this approach than conventional statistics. However, in the same vein, there are numerous questions for which there is no alternative but to use group statistics, whatever the flavour is conventional null-hypothesis significance testing, Bayesian hypothesis testing (Dienes, 2014; Rouder et al., 2009; Wagenmakers, 2007; Wagenmakers et al., 2018), or effect size estimation (Cumming, 2014). If the goal of the research is to have a precise knowledge of the magnitude of the effect, the latter is probably the best option. Often this will necessitate very large samples (de Winter et al., 2016). Similarly, even if the exact population effect size is not crucial, but an estimate is required to interpret the findings, then conventional group statistics are necessary. Naturally, if individual subjects only contribute singular measures (such as a questionnaire score, a behavioural performance level, or a neurophysiological amplitude, etc.) only the conventional group analysis is possible. The choice of statistical approach therefore depends on the research question and the experimental design. We hope that our exposition of the small-N approach will encourage researchers to consider how they could apply it to answer their research questions instead of mindlessly relying on conventional group statistics.

How should a researcher decide whether to use the small-N approach or rely on conventional group statistics? Testing the universality of an effect must necessitate a principled reason why universality is *plausible*. The examples we have discussed thus far, like the oblique effect, are based on scenarios where we have a reasonable idea of the ground truth. The small-N approach is only suitable in situations where the hypothesised effect is relatively “pure”, so that it can be isolated in each subject. Examples could be experiments using a classical prime or attentional cue that is predicted to speed up behavioural responses on a subsequent task. If we posit that this reflects the general function of human brains/minds, a sufficiently sensitive experiment should reveal this is in most subjects. In contrast, other scenarios, such as the embattled social priming studies, are unlikely to be testable using small samples. One famous study reported that subjects, who were primed to think of the concept of old age, would walk out of the lab more slowly (Bargh et al., 1996), a finding that has since been called into question (Doyen et al., 2012). The small-N framework inherently calls for a within-subject design and entails testing each subject over many trials. In the Bargh study, both these design changes could have resulted in the subject guessing the nature of the experiment and produced other long-term confounds like fatigue (not to mention making the experiment impractical to conduct). Most critically, a complex behaviour like how long it takes a subject to walk out the lab must involve a mixture of multiple cognitive and biological factors, ranging from their current emotional state to their physical fitness. This means inter-individual variability is large, and so even if the effect exists on average, it will not universally manifest in each person. By introducing our framework for small sample research, we implicitly encourage researchers to pay more attention to whether their statistical approach aligns with their research question and design their studies accordingly.

### Underlying assumptions

All statistical procedures are based on assumptions. The small-N approach is no exception. Since it is fundamentally based on binomial probabilities, it inherits the assumptions underlying binomial probabilities: the individual subjects must be independent of one another. Any relationship between individuals in the sample could create spurious findings; for example, if subjects are genetically related and the effect under investigation is partly heritable, the odds that all subjects show the same result are inflated. It is therefore important that the small sample captures a reasonable demographic breadth. Psychophysical studies should probably include a relatively large proportion of naïve subjects rather than only testing authors and their lab members who may be overtrained and familiar with the hypothesis (although as a counterpoint, note our example from the literature above). Inclusion and exclusion criteria should further ensure that the sample is indeed typical: when testing whether humans have an appendix, the study might exclude individuals with a history of abdominal surgery. The severity of this concern will depend on the exact nature of the research.

Another assumption underlying binomial tests is that the probability of success is equal for all observations. In our framework, this entails that the sensitivity is the same for all subjects, which may not hold in situations where there are individual differences in the effect. It is therefore advisable to assume the minimum sensitivity applies to all observations. Ideally, researchers should design their study in such a way that the sensitivity of individual tests is based on the *minimal effect of interest* (e.g., the smallest difference between psychometric curves considered meaningful). However, even in the absence of exact knowledge of sensitivity we can determine the strength of evidence by plotting the full sensitivity-evidence curve. This reveals the minimal sensitivity theoretically required for a conclusive result, given the observed results.

Importantly, no statistical test can replace theory. Formulating a clear hypothesis to test your research question is crucial. The small-N approach tests for the universality of the effect. It is the researcher’s responsibility to ensure whether such a hypothesis is realistic. If this is unlikely, or it is impossible to devise a sensitive test at the level of individuals, then the small-N framework is not the appropriate tool. But again, the same caveat applies to all other statistical models: using the wrong procedure can yield spurious inferences. Therefore, any use of the small-N test (or any other statistical method) should entail a justification why it is applicable.

Beyond maximising individual sensitivity, the small-N approach also requires practical decisions on experimental design. By setting a common likelihood for H_1_ for all subjects, the small-N test implicitly assumes that experimental conditions for individual subjects are closely matched. Differences in conditions would lead to variable sensitivity, which diminishes the overall sensitivity for H_1_. However, even with a perfect experimental design, external nuisance factors (such as a subject’s cognitive or physical state, or even the weather) could affect the sensitivity of individual tests. So do genuine individual differences between subjects, as a weaker effect requires greater sensitivity to detect. The good news is that less-than-perfect sensitivity can in fact *strengthen* the evidence for a universal effect. As the brown curves in Figure 2 illustrate, when not all subjects reach significance (e.g., 4 out of 6), the evidence ε is greater at intermediate sensitivity levels than at near-perfect levels: since the likelihood of detecting individual effects is well below 1, we should expect that not all subjects are significant.

### Statistical philosophies

Purists in statistical philosophy might balk at our mixture of frequentist statistical tests at the first (individual subject) stage and a likelihood approach at the second stage aggregating the evidence. However, we disagree with this objection on pragmatic terms. For one thing, our approach is extremely versatile. While most examples used here involve frequentist statistics at the first stage, this is not required. Our approach simply quantifies the number of subjects/tests successfully confirming a hypothesis. This can take many forms that do not involve formal significance tests; for example, a success could simply be based a performance criterion or some other numerical threshold. This criterion should of course be chosen to maximise sensitivity and minimise false positives. That said, in many situations one could construct a pure likelihood approach to reach similar conclusions as we do with our approach. For example, one study used a Bayesian hierarchical model to quantify the evidence of highly consistent findings like the Stroop effect (Rouder & Haaf, 2021). More generally, instead of individual significance tests one could use likelihood tests at the first stage. Quantifying the evidence would then simply involve multiplying the individual likelihoods. This approach is more sensitive because it does not binarise the individual tests. However, it necessitates that a likelihood ratio can be easily calculated for individual tests. In contrast, the small-N approach we describe here is easily understood without extensive knowledge of statistics and thus suitable to a wide user base.

### Concluding remarks

It should also go without saying that the small-N approach is not an excuse for lacking methodological rigour. In writing this article we in fact aim to enhance the rigour of studies using small samples. For the first time, this framework allows researchers to set predefined levels of evidence necessary to make inferences from their small-N data. This could also be useful for replication efforts. Because the small-N approach critically depends on the sensitivity of individual tests, it focusses researchers on maximising sensitivity. The approach also directs attention to other aspects of the experimental process, such as clearly defining exclusion criteria or pre-processing parameters. Despite their advantages, small-N studies are obviously not immune from the adverse effects of flexible methods. Experimental procedures and analysis methods should normally be determined – and ideally preregistered – prior to data collection. Thus far, without any way to determine evidence thresholds small sample studies have been automatically precluded from enjoying methodological advances like *Registered Reports* in many journals (Chambers, 2013; Chambers et al., 2015). Our framework now offers researchers greater control of biases arising from methodological flexibility and boost the credibility of scientific research based on small samples. Lastly, like all other statistical methods, the small-N test should not be applied mindlessly. However, by design the framework encourages deeper considerations of the experimental design to maximise sensitivity and ensure the independence of individual subjects. We therefore hope that the test provokes more discussion about those aspects, both in the planning stage and in peer reviews of completed studies.

## Data availability

Functions for MATLAB, R, and an Excel spreadsheet for calculating the Small-N evidence ratio, as well as analysis code and data from the example oblique effect experiment are publicly available at: https://osf.io/bmzc3 (Schwarzkopf & Huang, 2023a). We also published an earlier version of this article as a preprint (Schwarzkopf & Huang, 2023b).

## Acknowledgements

We thank Zoltan Dienes, Benjamin de Haas, and Will Harrison, as well as Klaus Fiedler and two anonymous reviewers for constructive feedback on this manuscript. We also thank the editors and reviewers at multiple journals who inspired us to write this article by insisting that only ever-larger samples can yield meaningful results.

## Supplementary Material

### Methods

#### Participants

Twenty participants (age range: 18-48 years, 15 female) with normal or corrected-to-normal visual acuity participated in the study. All gave written informed consent. The University of Auckland Human Participants Ethics Committee approved experimental procedures. Three participants were excluded from analysis because they failed to comply with task instructions.

#### Stimuli

Stimuli were generated using MATLAB (MathWorks, Inc.), and the experiment was presented using the online platform Testable (https://www.testable.org). The stimuli were Gabor patterns with different orientations, sinusoidal gratings (wavelength: 1.5°) at maximum contrast convolved with a two-dimensional Gaussian (standard deviation: 1°). We used three main conditions with orientations centred on 45°, 90°, or 135°. The orientation of test patterns could differ from these main orientations by 0°, ±5°, ±10°, ±15°, or ±20°. All stimuli were presented on a uniform grey screen.

#### Procedure

Participants sat in front of a computer in a small testing cubicle, at a distance of 75 cm from the screen (model: Dell OptiPlex 5260 aio series, size: 26.8 cm by 47.6 cm, 21.5 inch, resolution: 1920 x 1080 pixels). There were 27 unique trial types: nine orientation offsets for each of the three main orientations. Each trial type was presented 20 times in the experiment in interleaved order. Therefore, there were 540 trials in the actual experiment. In addition, 12 trials at the beginning of the task were practice trials that were not used for analysis.

Each trial contained two stimuli. The first grating was the reference, presented for 500 ms. This could be one of the three main orientations. This was followed by a 300 ms inter-stimulus interval during which only a grey screen and the fixation cross were presented. The second grating appeared on the screen until the participant responded. It could be tilted anticlockwise, clockwise, or remain unchanged relative to the first grating (Figure S1). Participants indicated via the keyboard left or right arrow keys whether the second grating was tilted anticlockwise or clockwise, respectively. They were instructed to respond as fast and as accurately as possible. If they were unsure about the change between the two stimuli, they must nevertheless press one key to continue the task. Throughout the run, participants were asked to fixate on a small, black cross in the middle of the screen. After the participant’s response, a blank screen was presented for a 500 ms inter-trial interval.

During the 12 initial practice trials, participants could only choose the correct answer to move on to the next trial. The stimuli in the practice were easy to discriminate, including only orientation offsets of ±10° and ±20° for each of the three main orientations. Most participants could select the correct answer easily.

**Figure S1.**
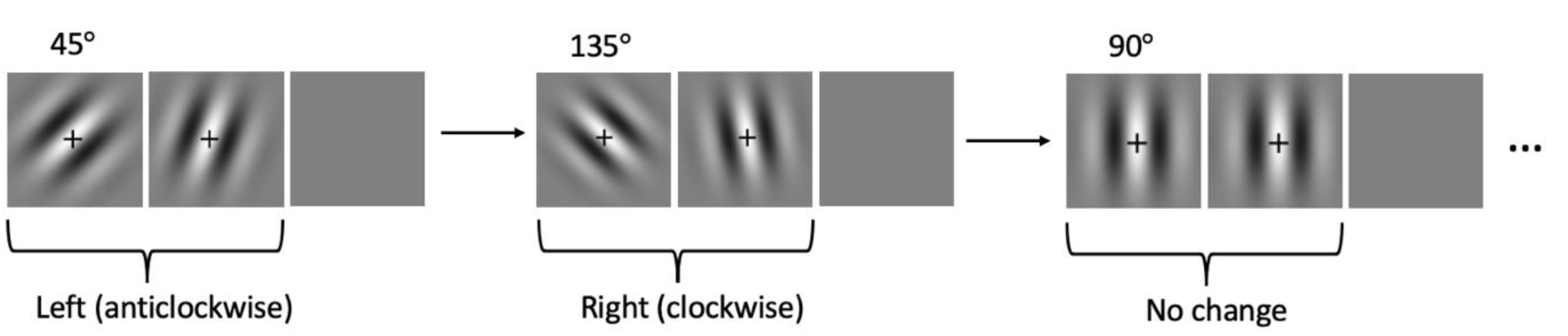
Experimental design. *Trial sequence.* The second (test) grating was either tilted anticlockwise or clockwise or was unchanged relative to the first (reference) grating. There were three main orientations around which the test grating could vary. The oblique effect is characterised by superior performance around cardinal (in this case, vertical) orientations. Note that a 500 ms inter-trial interval followed the observer’s response. There was also a 300 ms inter-stimulus interval between the two gratings in each trial, not shown here. The second grating stayed on the screen until the observer responded.

#### Data analysis

We then determined the participant’s orientation discrimination performance for each of the three main orientations. For each orientation offset, we calculated the proportion of trials that they chose the test stimulus as appearing more anticlockwise. Then we fit a cumulative Gaussian psychometric function to these data using a linear mean squares approach. The bandwidth (σ) of this Gaussian determines the slope of the curve. Narrower bandwidth corresponds to steeper slopes, meaning better performance (Figure S2A). We calculated two contrasts of interest. For determining the oblique effect (H_1_) we subtracted the bandwidth for 90° main orientation from that for 45°. To estimate an assumed null effect (H_0_), we subtracted the bandwidth for 135° from that for 45°.

We also determined the significance of these contrasts for each observer to use these results in our small-N approach. To this end, we bootstrapped the psychometric curve fits. We resampled the choices for each orientation offset with replacement, recalculated the choice proportions, and refit the psychometric curves 10,000 times. We then determined a one-tailed significance value by calculating the proportion of resamples for which the bootstrapped distribution of the two contrasts were ≤0.

## Results

Psychometric curves for three representative observers are shown in Figure S2A. It is immediately apparent that the curve for 90° main orientation (red functions) has a steeper slope in all observers, that is a narrower bandwidth. We plot the contrasts between bandwidths for H_1_ (45°-90°) and H_0_ (45°-135°) in Figure S2B. All observers showed positive contrasts for H_1_ as expected, while contrasts for H_0_ ranged between positive and negative but most were in fact negative. Only one observer failed to reach significance on individual fits for H_1_ and only one was significant for H_0_.

**Figure S2.**
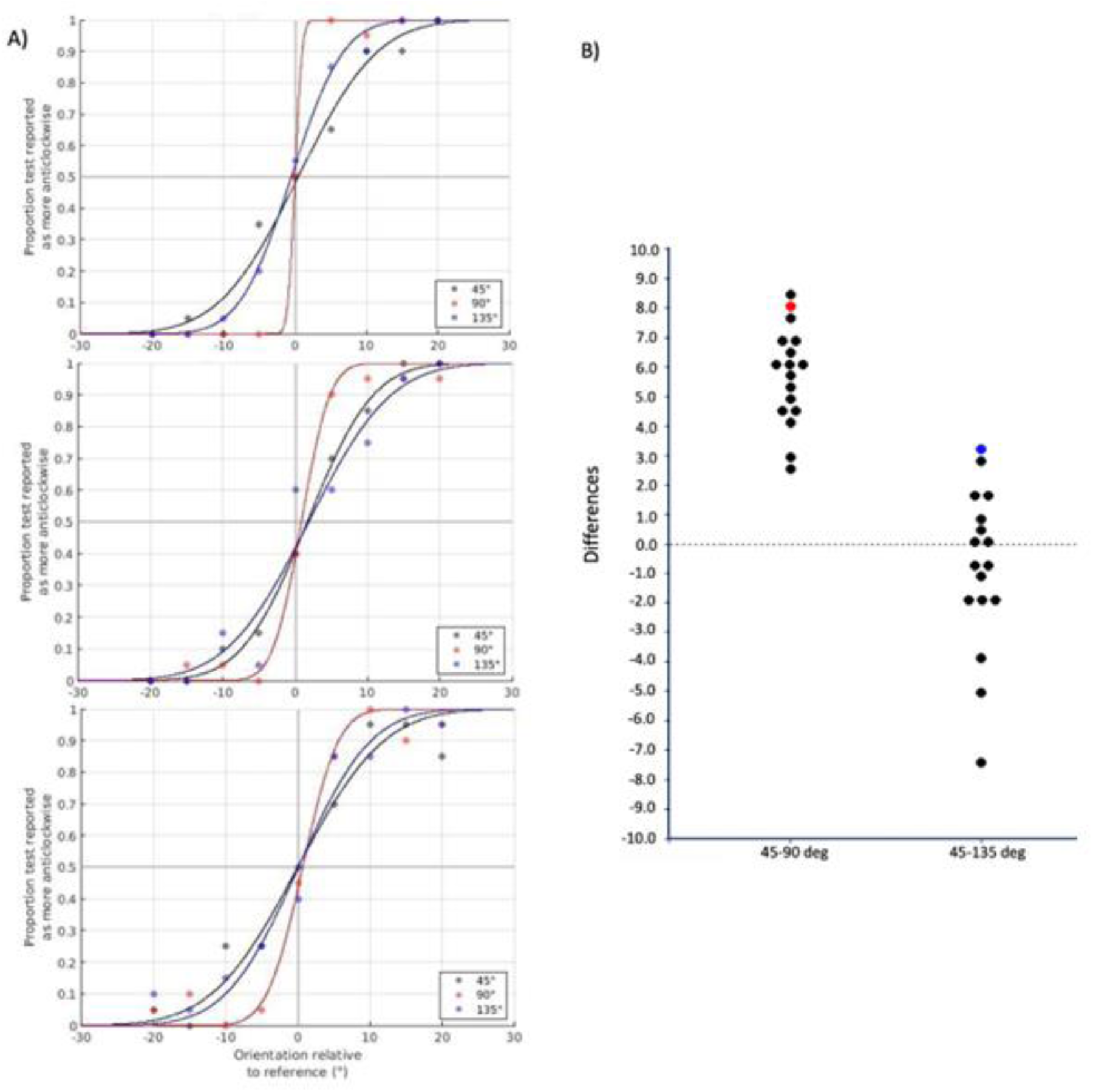
Oblique effect results. **A)**. Psychometric curves from three representative observers. The proportion of trials the test grating was seen as more anticlockwise is plotted against the orientation offset, separately for the three main orientations. Black: 45°, Red: 90°, Blue: 135°. **B)**. Contrasts across participants. Each dot denotes the difference in bandwidth between the conditions shown. The red dot indicates the observer who failed to reach significance for the contrast 45-90°. Conversely, the blue dot indicates the observer who showed a significant difference for the contrast 45-135°.

## Footnotes

1 For simplicity, throughout this article we refer exclusively to ‘subjects’. We mean this to include all individual tests that could be envisioned in a small-N design, ranging from human participants or animals in an experiment to other cases, such as tissue samples, testing sites, etc. When discussing examples from psychophysics studies, we instead use the term ‘observer’ to clarify that this refers to that specific situation.

